# Protein painting reveals pervasive remodeling of conserved proteostasis machinery in response to pharmacological stimuli

**DOI:** 10.1101/2022.05.14.491969

**Authors:** Dezerae Cox, Angelique R. Ormsby, Gavin E. Reid, Danny M. Hatters

**Affiliations:** Department of Biochemistry and Pharmacology, Bio21 Molecular Science and Biotechnology Institute, The University of Melbourne, Parkville, VIC 3010, Australia; Department of Chemistry, University of Cambridge, Cambridge CB2 1EW, United Kingdom; School of Chemistry, The University of Melbourne, Parkville, VIC 3010, Australia

**Keywords:** Proteostasis, proteasome, molecular chaperone, proteomics, interactome

## Abstract

Accurate spatio-temporal organization of the proteome is essential for cellular homeostasis. However, a detailed mechanistic understanding of this organization and how it is altered in response to external stimuli in the intact cellular environment is as-yet unrealized. To address this need, ‘protein painting’ methods have emerged as a way to gain insight into the conformational status of proteins within cells at the proteome-wide scale. For example, tetraphenylethene maleimide (TPE-MI) has previously been used to quantify the engagement of quality control machinery with client proteins in cell lysates. Here, we showcase the ability of TPE-MI to additionally reveal proteome network remodeling in whole cells in response to a cohort of commonly used pharmacological stimuli of varying specificity. We report specific, albeit heterogeneous, responses to individual stimuli that coalesce on a conserved set of core cellular machineries. This work expands our understanding of proteome conformational remodeling in response to cellular stimuli, and provides a blueprint for assessing how these conformational changes may contribute to disorders characterized by proteostasis imbalance.

## Introduction

Precise spatio-temporal regulation of the proteome is essential for cellular homeostasis. This occurs at various levels, from the folding of individual protein domains, to binary protein-protein interactions, and the assembly of multi-protein macromolecular machines. The result is the culmination of protein networks that drive biological functions. The protein networks are usually interconnected with other networks to mediate their regulation or to direct sequential functions (such as signaling pathways). A mechanistic understanding of cellular (dys)function in health and disease requires detailed knowledge of this organization and how it is altered in response to external stimuli.

An outstanding challenge has been the capacity to quantitatively assess the macromolecular organization of individual proteins in cells at the proteome-wide scale. After decades of dedicated examination, the folding and stability characteristics of many individual proteins are well understood *in vitro*. Thus far, high-throughput approaches to quantify protein conformation have included proteomic variations of these *in vitro* methodologies, relying on either the accessibility of protein regions to non-specific proteases (Liu & Fitzgerald, 2016; Schopper *et al*, 2017), thermal aggregation-based methods (Leuenberger *et al*, 2017; Tan *et al*, 2018), or basal protein solubility (Sui *et al*, 2020; Wallace *et al*, 2015). A map of all possible binary protein-protein interactions has also been aggressively pursued in yeast (Luck *et al*, 2020). However, these methods are all limited by the need to assess proteins either outside the cellular environment (i.e., *ex vivo*, post lysis) or under conditions of altered protein expression, and often cannot probe subtle changes in proteome organization within the undisturbed cellular context.

Protein painting methods have emerged as a way to gather conformational insight within intact cells at the proteome-wide scale. Recently, we described the application of one such method based on a fluorogenic dye, tetraphenylethene maleimide (TPE-MI). TPE-MI reacts with exposed cysteine free thiols that are otherwise buried in the folded state. Free cysteine thiols are the least surface-exposed residue of all amino acids in globular proteins, and thus provide an excellent target for examining protein conformation (Marino & Gladyshev, 2010). We have previously reported the ability of this probe to provide a snapshot of proteome conformation in live cells (Chen *et al*, 2017), and more recently explored aspects of proteome organization in response to denaturation in cell lysate (Cox *et al*, 2022). Here, we extend this methodology to explore remodeling of proteome networks in response to a cohort of commonly used pharmacological stimuli. We detect specific, heterogeneous responses to individual stimuli, however these changes in proteome organization coalesce on a conserved set of core cellular machineries.

## Results

### Pharmacological stimuli induce changes in proteome conformation

We selected a cohort of pharmacological stimuli that modulate different aspects of cellular homeostasis, and which act with varying degrees of specificity. The mechanism of action and relative specificity for these compounds is summarized in Table 1. Two compounds were selected that have potent, reversible, and specific targets: the synthetic peptide aldehyde MG132 (Z-Leu-Leu-Leu-al; (Goldberg, 2012)), which inhibits proteolytic activity of the proteasome by specific interaction with the ß5 (and at high concentration, the ß1) subunits of the 20 S proteasome; and the small molecule inhibitor VER155008, which inhibits the chaperone activity of Hsp70 family proteins by binding to the ATPase domain (Schlecht *et al*, 2013; Williamson *et al*, 2009). The third compound, staurosporine, was selected as a prototypical ATP-competitive kinase inhibitor that binds non-selectively to kinases with high affinity (Karaman *et al*, 2008). Thus, while still characterized by a specific mechanism of action, the target range of staurosporine is comparatively large. The final two stimuli were selected as having well-characterized broad-spectrum activities. Celastrol is often used as an inducer of the heat-shock response due to its ability to activate HSF1 (Westerheide *et al*, 2004), however it also has a range of off-target effects including inhibition of the proteasome and HSP90 chaperones (Yang et al, 2006; Zhang et al, 2009). Novobiocin is an antibiotic for Gram-positive pathogens that inhibits bacterial DNA gyrase by binding the ATP-binding site in the ATPase subunit (Neckers *et al*, 2018; Marcu *et al*, 2000). In mammalian cells it has a lower level affinity to the C-terminal nucleotide binding pocket of Hsp90, inhibiting its chaperone activity with an IC_50_ of ∼700 µM (Donnelly & Blagg, 2008; Marcu *et al*, 2000; Burlison *et al*, 2006). Unlike other modifiers that target the N-terminal domain of Hsp90, novobiocin does not induce a heat shock response (Terracciano *et al*, 2018). However, as a low affinity Hsp90 inhibitor it would be expected to have high levels of off-target activity (Burke *et al*, 1979; Edenberg, 1980).

**Table 1:** Pharmacological stimuli. Treatments were diluted into fresh culture media before incubation at 37 °C. Compounds are categorized as poor (+), moderate (++) and high (+++) specificity according to the scope of target and reported range of off-target effects.

| Compound | Specificity | Source | Vehicle | Concentration | Incubation time (h) |
| --- | --- | --- | --- | --- | --- |
| MG132 | +++ | Sigma #C2211-5MG | DMSO | 10 µM | 18 h |
| VER155088 | +++ | Sigma #SML0271 | DMSO | 20 µM | 18 h |
| Staurosporine | ++ | AbCam #ab146588 | DMSO | 500 nM | 2 h |
| Celastrol | + | Sigma #0869 | DMSO | 5 µM | 18 h |
| Novobiocin | + | Sigma #N1628 | mQ | 800 µM | 6 h |

To explore proteome remodeling in response to these diverse pharmacological stimuli, we deployed the cysteine-reactive fluorogenic probe TPE-MI in a model neuron-like system, Neuro-2a cells. Cells treated with each compound were labeled with TPE-MI, then analyzed via flow cytometry (Fig. 1A). The median fluorescence of the main cell population (derived as described in (Chen *et al*, 2017)) demonstrated significant increases in TPE-MI fluorescence in cells treated with each of the compounds (Fig. 1B). This net increase in the global exposure of buried thiol residues suggests significant proteome rearrangement.

**Figure 1:**
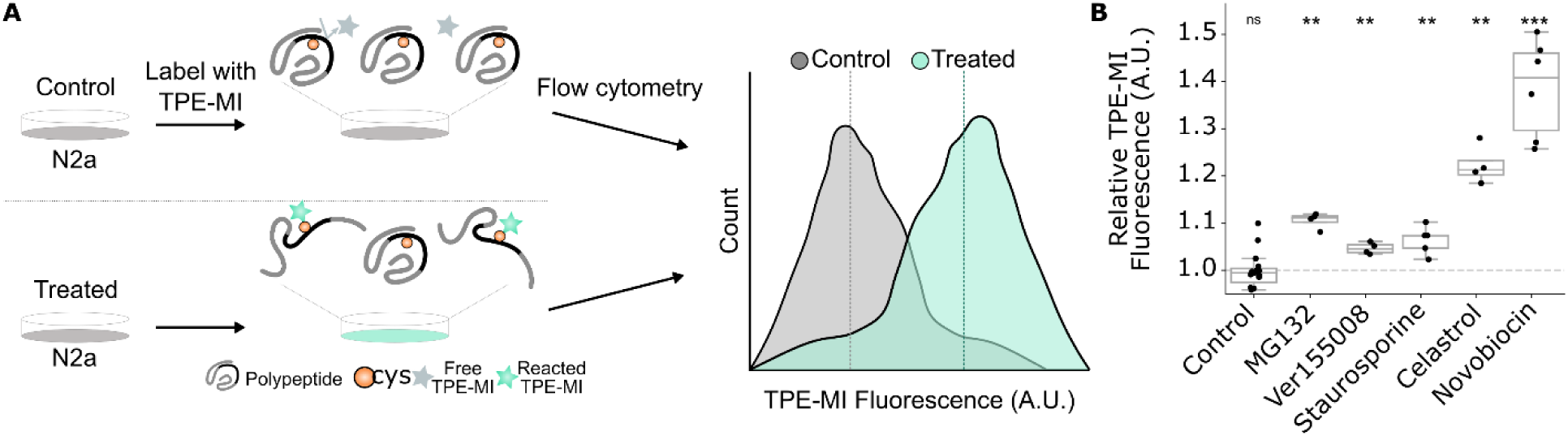
Pharmacological stimuli result in net increases in TPE-MI reactivity of cellular proteins. (A) Method schematic for quantifying global proteome conformation. Neuro-2a cells were treated with MG132, VER155008, staurosporine, celastrol or novobiocin before labelling with TPE-MI. Cells were then analyzed via flow cytometry. (B) Median TPE-MI fluorescence measured at 740 nm, normalized to the vehicle-treated control population. Shown are boxplots overlayed with individual datapoints of at least four replicates (dots), * p<0.05, **p<0.01, *** p<0.001 according to one-sample t-test against hypothetical mean of 1.

### Proteasome inhibition remodels protein complexes associated with apoptosis

We next assessed the contribution of individual proteins to global changes in proteome organization using proteomic analysis (Chen *et al*, 2017; Cox *et al*, 2022); Fig. 2). Briefly, SILAC-labelled Neuro-2a cells treated with either the vehicle control (light) or the compound of interest (heavy) were labelled with TPE-MI then subjected to LC-MS/MS. Changes in the reactivity of cysteine thiols in individual proteins were quantified using the ratio of cysteine-containing peptides between compound- and vehicle-treated cells, after correcting for any change in total per-protein abundance according to the non-cysteine containing peptides (Corrected cys ratio; see Methods for details). The resultant change in reactivity of cysteine-containing peptides constitutes a reporter of changes in protein conformation. Finally, p-value weighted scaling and data-driven thresholds were applied (Cox *et al*, 2022) such that changes in corrected cysteine ratios outside the control thresholds were considered biologically of interest.

**Figure 2:**
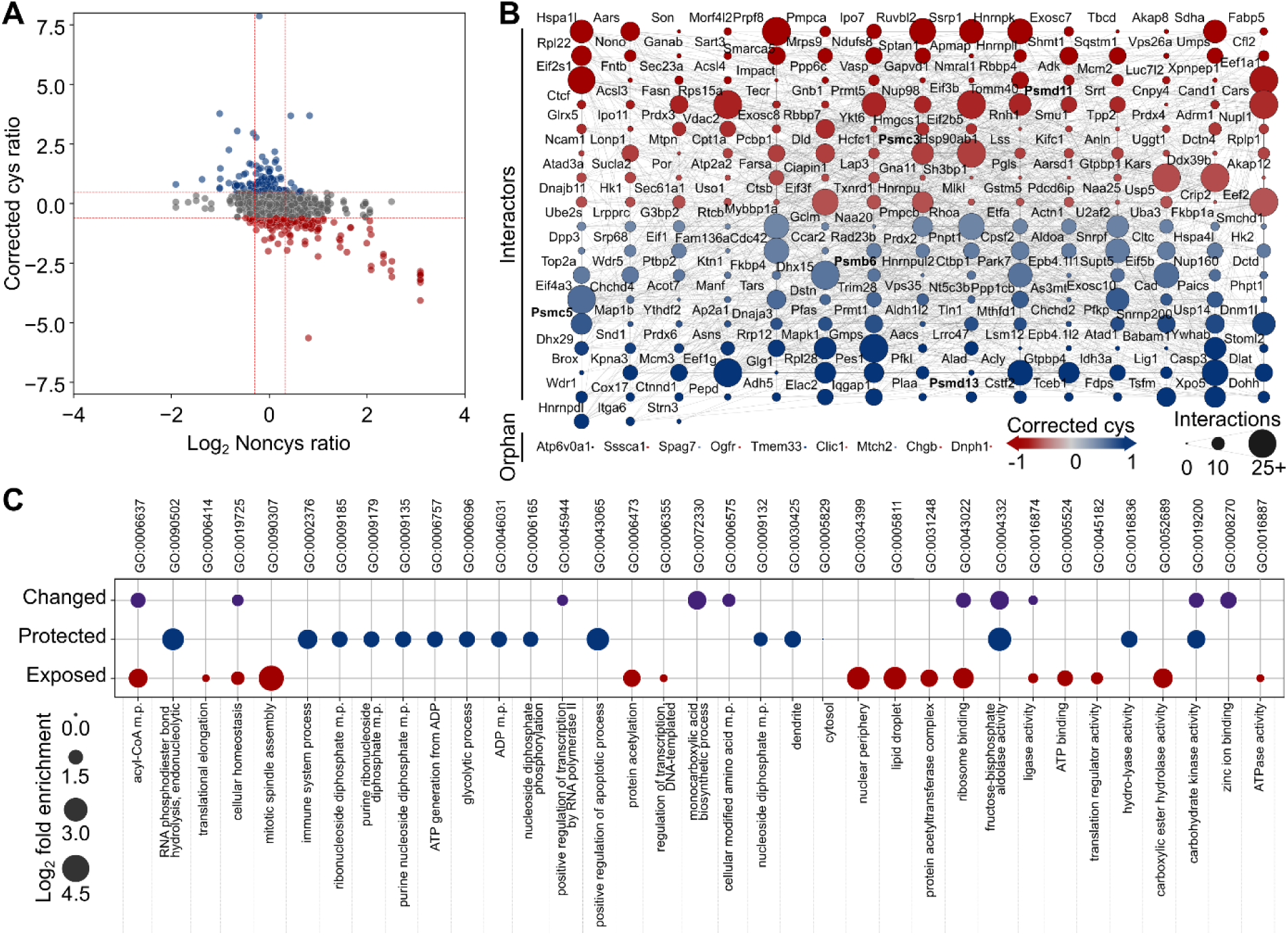
Conformational changes due to proteasome inhibition reflect cellular responses to MG132. SILAC-labelled Neuro-2a cells were treated with a pharmacological stimulus i.e., MG132 (heavy) or vehicle control (light), then labelled with TPE-MI and prepared for proteome analysis using LC-MS/MS. Peptide quantitation yielded the relative abundance of noncys-containing peptides (measure of abundance) and cys-containing peptides (measure of conformational status). (A) Representative scatterplot for processed noncys- and cys-peptide abundances in the presence of proteasome inhibitor MG132. Thresholds (red dotted lines) determined based on control dataset are shown, outside which cysteine-containing peptides were considered more exposed (red) or more protected (blue). (B) Protein interaction network for changed peptides derived from B. Protein nodes are colored according to maximum corrected cys ratio and edges (lines) connect proteins with known interaction (STRINGdb v 11.0, medium confidence score > 0.4). Protein nodes are sized according to the number of interactions within the network. (C) Significantly enriched gene ontology terms (p < 0.05) for all proteins which changed reactivity (purple); or more specifically became protected (blue) or exposed (red) in B. Enrichment terms filtered to minimize hierarchical redundancy (PantherGOSlim v 16.0).

The cysteine thiol reactivity profile under conditions of MG132-mediated proteasome inhibition is shown in Fig. 2A (results for the remaining treatments are summarized in Fig. S1). Of the 2880 cysteine-containing peptides quantified, representing 1163 proteins, 306 were seen to change reactivity. As a whole, these proteins formed a densely connected protein-protein interaction network (Fig. 2B; STRINGdb enrichment test, p<0.0001), and were enriched for machinery associated with regulating biological quality (GO:0065008) and cellular homeostasis (GO:0019725) (Fig. 2C). Of these, 164 cysteine thiols increased in reactivity; the corresponding 129 proteins were enriched for machinery associated with biosynthesis and protein production (Fig. 2C), including ribosome binding (GO:0043022), regulation of transcription DNA-templated (GO:0006355) and translation (GO:0006412). In contrast, 142 cysteine thiols, representing 131 proteins, were seen to decrease in reactivity. We have previously associated this protection phenomenon with functional complex remodeling (Cox *et al*, 2022). Proteins exhibiting protection in response to MG132-mediated proteasome inhibition included three proteasome subunits (PSMC5, PSMB6 and PSMD13), and were enriched with machinery associated with positive regulation of apoptotic processes (GO:0043065) and cell death (GO:0010942) (Fig. 2C). Broadly, this is consistent with cellular phenotypes previously observed in response to proteasome inhibition with MG132, including reconfiguration of transcription and translation (Cowan & Morley, 2004; Heine *et al*, 2008) and induction of apoptosis (Yuan *et al*, 2008).

### Proteome organization is fine-tuned across core cellular activities in response to pharmacological stimuli

After applying the same methodology to the remaining stimuli, we compiled a list of 646 proteins which were quantified in all treatments and whose TPE-MI reactivity was altered by at least one stimulus (referred to hereon as the comparison protein set). Individual proteins were poorly conserved in their response to each treatment, such that less than 1% of these proteins were seen to have altered reactivity as a result of all five compounds, which is indicative of the highly distinct mechanisms by which the stimuli act on cells (Fig. 3A). 50% of proteins featured altered reactivity specific to a single treatment. In contrast, almost 10% of the comparison proteins were found to have modified reactivity that were common to the celastrol, MG132 and novobiocin stimuli. This is consistent with the fact that the combined targeted and off-target effects of these stimuli intersect. This supports our ability to ascribe the response of individual proteins to distinct stimuli and demonstrates that we are not merely measuring generic changes in a subset of proteins in response to any miscellaneous treatment.

**Figure 3:**
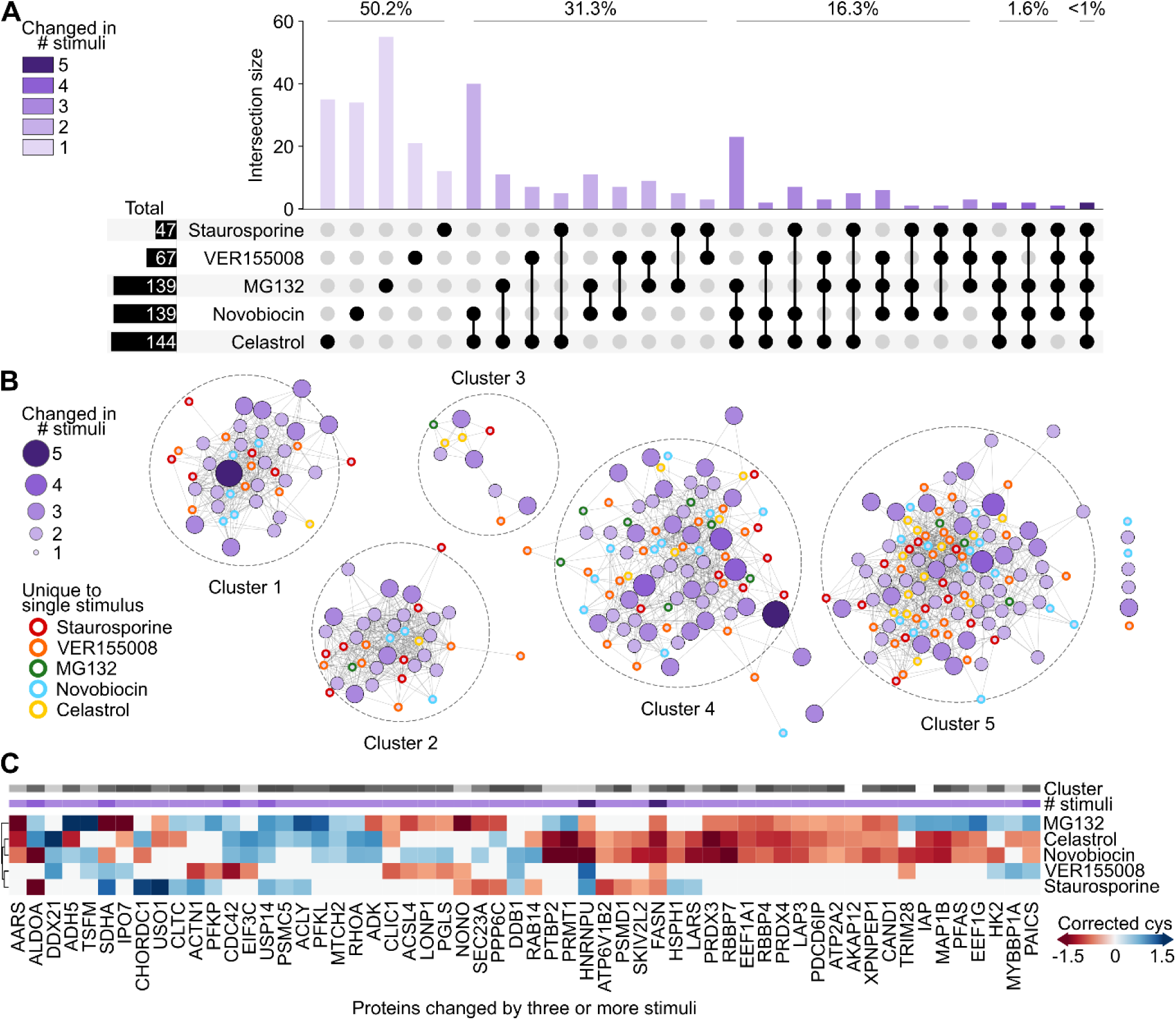
Conformational remodeling is poorly conserved in individual proteins in response to diverse stimuli. Proteins which were quantified in each of the five stimulus datasets, and which changed reactivity in at least one treatment, were collected as the comparison protein set. (A) UpSet intersection plot of conformational change among comparison proteins. Individual proteins appear once within the column bar graph, according to the combination of treatments in which they were seen to change conformation (irrespective of reactivity direction). The proportion of proteins associated with change in 5, 4, 3, 2, or 1 treatment are indicated above the corresponding bars. (B) Protein interaction network for comparison proteins. Protein nodes are colored and sized according to their degree of commonality across treatments in A, and for first-degree proteins the corresponding treatment is indicated by border color. Nodes were arranged organically following clustering with Girvan-Newman community detection algorithm, and edges (lines) connect proteins with known interactions within each cluster (STRINGdb v 11.0, medium confidence score > 0.4). (C) Heatmap for maximum change in cys reactivity among comparison proteins with degree > 3 from A. Both degree (# compounds) and cluster number are indicated for each protein above the heatmap, and dendrogram shows the result of agglomerative clustering of the filtered comparison proteins in the treatment dimension.

We next sought to assess features of the comparison proteins. The protein-protein interaction network among these proteins was significantly more connected than would be expected by chance among a group of equivalent size (STRINGdb enrichment test, p<0.0001). As with the proteasome inhibition experiment described above, this suggests functional groupings within the comparison proteins. To further investigate this, we clustered the protein-protein interaction map using the Girvan-Newman fast greedy algorithm for community detection (Newman & Girvan, 2004). This produced five major clusters of densely connected proteins (Fig. 3B), and seven additional ‘orphan’ proteins. Additional gene ontology analysis of proteins in each cluster revealed enrichment patterns reminiscent of core cellular activity hubs; namely, transcription (cluster 1), translation (cluster 2), intracellular trafficking (cluster 3), enzymatic activity and biosynthesis (cluster 4) and protein synthesis and degradation (cluster 5) (Fig. S2). There was no discernable pattern of commonality or response to individual compounds among the clusters; all five clusters contained proteins whose conservation ranged from two to at least four treatments. In addition, those proteins whose conformational change was unique to a single compound were similarly spread across all five clusters.

As well as the binary measure of conformational change, we also considered the maximum change in cysteine reactivity per protein associated with individual stimuli (Fig. 3C; Fig. S3). We observed an additional layer of heterogeneity in response to individual stimuli; i.e., proteins often became exposed in response to one compound but protected as a result of another. This was reminiscent of our previous study, in which heterogeneous changes in proteome solubility resulted from proteostasis imbalance (Sui *et al*, 2020). However, despite identifying more than 90 % identical proteins, there was no significant correlation between reactivity and solubility changes in any of the matched treatments (MG132, novobiocin and VER155008; Fig. S4). This is consistent with the ability of TPE-MI to measure subtle changes in proteome organization when compared to an aggregation based methodology (Cox *et al*, 2022).

We further quantified this heterogeneity by measuring the correlation between comparison proteins according to several grouping features. We found no correlation between the reactivity response of a protein and its degree of conservation across treatments (spearman’s correlation coefficient = -0.092 for all datapoints; Fig. S5). Similarly, there was no significant correlation among proteins associated with individual KEGG pathways (a collection of pathway maps containing molecular interactions, reactions, and relation networks responsible for cellular metabolism, structure, and information processing). However, the average magnitude of correlation (R_s_) for each protein with partners inside the same functional cluster was significantly different to those outside the cluster (Fig. 4A; t-test, p = 0.032). Similarly, reactivity changes were significantly more correlated among protein interaction partners (Fig. 4B; t-test, p < 0.001).

**Figure 4:**
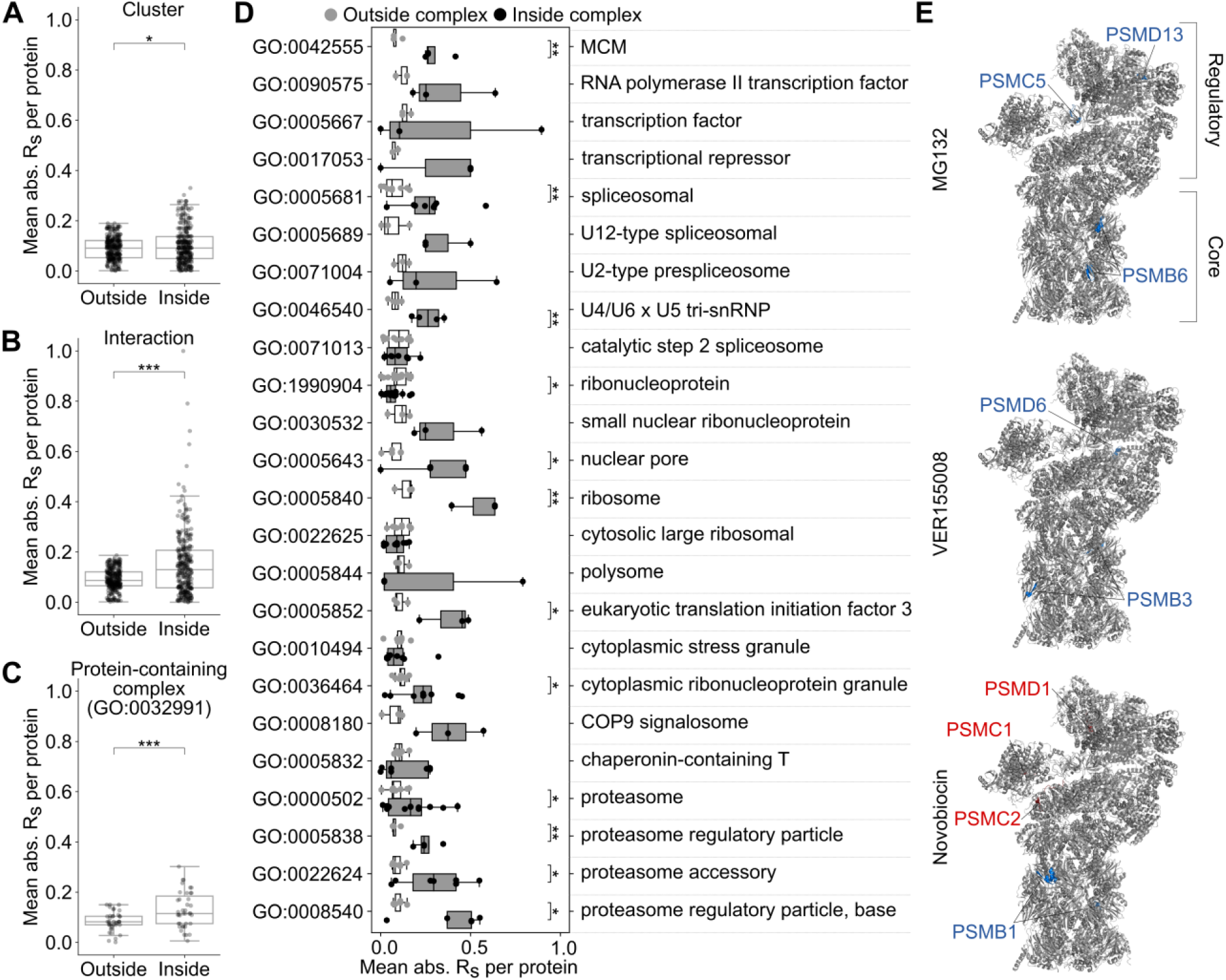
Unique conformational remodeling of macromolecular complexes is specifically associated with different stimuli. Correlation comparison for (A) functional clusters presented in Fig. 3, (B) interacting proteins, and (C-D) proteins annotated with gene ontology terms associated with macromolecular complexes. The pairwise correlation strength was calculated for all comparison proteins then binned according to whether one (outside) or both (inside) proteins identified with the feature of interest. The mean for each protein in both bins was then calculated and the resultant inside and outside values were compared according to a two-tailed t-test. * p<0.05, **p<0.01, *** p<0.001. (E) Human 26S proteasome ribbon structure adapted from PDB: 6MSB. Variations of the structure are presented for each of the MG132, VER155008 and novobiocin conditions, whereby individual cysteine-containing peptides are colored according to their increase (red) or decrease (blue) in reactivity. The corresponding protein subunits are labelled with an equivalent color scheme.

### Conformational changes are consistent with remodeling of macromolecular complexes

In addition to the presence of a one-to-one interaction between comparison proteins, we wondered if proteins associated with specific macromolecular complexes may be similarly correlated. We first considered proteins annotated with the generic gene ontology term “protein-containing complex” (GO: 0032991) and found that these proteins were significantly more correlated with each other than with non-complex proteins (Fig. 4C; t-test, p < 0.001). Exploiting the hierarchical nature of gene ontology annotations, we then compared the average correlation among proteins for individual complexes which fall under the protein-containing complex umbrella (Fig. 4D). We identified several complexes for which reactivity changes were significantly correlated, including transcription, translation, and degradation machinery. This suggests at least some of the changes in reactivity we observe are a result of the assembly and/or disassembly of individual macromolecular complexes.

We explored this theory in the context of the 26S proteasome, components of which were among the significantly correlated complex terms (regulatory (GO:0008540) and accessory (GO:0022624) particles; Fig. 4D). The 26S proteasome is comprised of two subcomplexes: the catalytic core particle (so-called 20S proteasome) and one or two terminal activating regulatory particles (so-called 19S particles) (Bedford *et al*, 2010). The core particle forms an enclosed cavity where catalytic threonine residues contributed by PSMB5, PSMB6 and PSMB7, possessing chymotrypsin-like, caspase-like, and trypsin-like activity respectively, degrade substrate. The regulatory particles associate with the termini of the barrel-shaped core particle, where they recognize ubiquitylated client proteins and assist in their unfolding and translocation into the β-ringed catalytic chamber. Degradation by the 26S proteasome is ATP dependent and in most cases requires the presence of a ubiquitin chain on the substrate protein (Verma & Deshaies, 2000). However, the 20S core greatly outnumbers capped proteasomes in cells under basal conditions (Fabre *et al*, 2014, 2015; Tanahashi *et al*, 2000) and can degrade completely unfolded proteins in an ATP- and ubiquitin-independent manner (De Mot *et al*, 1999) in response to oxidative stress (Pickering & Davies, 2012).

The structure of the human 26S proteasome is well characterized and conserved in eukaryotes, providing a scaffold onto which we could map observed changes in reactivity for individual cysteine thiol containing peptides that resulted from different stimuli (Fig. 4E). Subunits located within the core 20S particle saw conserved protection as a result of MG132, VER1550088 and novobiocin. This is consistent with an increase in occupation of the catalytic chamber, which we anticipate as a response to the accumulation of unfolded proteins. Member subunits of the 19S regulatory particle (PSMC5, PSMD6 and PSMD13) were protected in the presence of MG132 and VER155008 (Fig. 4E). This may reflect increased assembly and engagement of the 19S regulatory particle with the surplus of core complexes to enhance specific ubiquitin-mediated degradation of unfolded proteins. In addition, PSMC5 and PSMD6 are located at the interface between the lid and base regions of the regulatory particle, undergoing key conformational changes that facilitate switching between substrate-free and substrate-bound states of the proteasome (Greene *et al*, 2019). In contrast, in the presence of novobiocin, three regulatory particle subunits (PSMC1, PSMC2 and PSMD1) became more reactive (Fig. 4E). This is consistent with evidence that HSP90 is required for *de novo* assembly of the 26S proteasome (Yamano *et al*, 2008), and that loss of HSP90 activity results in the disassembly of existing 26S proteasomes (Imai *et al*, 2003) followed by rapid dissociation of the regulatory particle components. Together, these examples demonstrate how seemingly heterogeneous changes in per-protein reactivity can reveal functional and specific remodeling of macromolecular complexes in response to different stimuli, even when the stimulus is expected to have a large extent of off-target activity.

## Discussion

We demonstrate here the application of TPE-MI to quantify conformational changes associated with a diverse range of pharmacological stimuli at the whole-cell and proteome-wide scales. Bulk measurements of increased TPE-MI reactivity associated with proteome unfolding were seen to mask subtle differences in reactivity at the per-protein level. Although per-protein changes were largely unique to a specific stimulus, these changes occurred in a conserved set of functional machinery which broadly match core cellular activities. These conserved hubs exhibited heterogeneous changes in response to different stimuli, hinting at finely tuned control of proteome conformation in response to stimuli that is commensurate with their degree of specificity.

Significant correlations among proteins known to interact enabled us to ascribe many of the observed changes in reactivity to the remodeling of protein-protein interactions, including within multi-subunit macromolecular complexes. The detailed structure-function information available for the human proteasome allowed us to rationalize the heterogenous changes observed for individual subunits of the 26S core and regulatory particles in response to different treatments. The ability to obtain mechanistic details for other macromolecular machines remains challenging. However, in addition to existing structural models, the advent of machine-learning approaches such as AlphaFold-multimer (Evans *et al*, 2021), which enables the prediction of protein-protein interaction interfaces from protein sequence alone, promises to assist in evaluating our observations of potential binding/unbinding events *in silico* to direct *in vitro* and *in vivo* validation efforts.

As Luck and colleagues observe (Luck *et al*, 2020), it remains infeasible to assemble a map of proteome organization in the context of the many thousands of physiological and pathological cellular contexts by systematically identifying endogenous protein-protein interactions (PPIs). However, the data reported here demonstrates the potential for protein painting technologies such as TPE-MI to provide detailed inventories of remodeling events that occur in response to stimuli within the intact cellular environment, and under conditions where complex changes are arising. There are two outstanding limitations of this method; namely, the inability to monitor proteins which don’t contain a free cysteine thiol residue, and the failure to adequately quantify some proteins across all treatment regimes which meant they were subsequently removed from the comparison dataset. Both of these will be readily addressed by combining TPE-MI with complementary protein painting strategies, for example lysine modification (Bamberger *et al*, 2020), and by leveraging ongoing advancements in data-independent acquisition and quantitation methodologies (Guan *et al*, 2020). Overall, this work expands our understanding of proteome conformational remodeling in response to cellular stimuli, and provides a blueprint with which to assess how these conformational changes may contribute to disorders characterized by proteostasis imbalance.

## Materials and Methods

### Materials

All materials used in this study were purchased from Sigma-Aldrich (St. Louis, MO, USA) unless otherwise indicated. The mouse neuroblastoma cell line Neuro-2a (N2a) was obtained from lab cultures originating from the American Type Culture Collection, and cultures were routinely screened for mycoplasma contamination. TPE-MI was a kind gift from Dr Yuning Hong (La Trobe University), and stocks prepared at 10 mM in DMSO were stored in the dark at 4 °C before use. All work was completed with protein low-bind plastics unless otherwise indicated.

### Cell culture

Neuro-2a cells were cultured in Dulbecco’s modified Eagle’s medium (DMEM; ThermoFischer Scientific) supplemented with 10% (v/v) fetal bovine serum (ThermoFischer Scientific) and 1 mM L-glutamine (ThermoFischer Scientific). In the case of isotopically labelled cultures (SILAC), cells were cultured in DMEM (Silantes) supplemented with either unlabeled (light) or ^13^C L-Lysine and ^13^C,^15^N L-Arginine, along with 10% (v/v) dialyzed fetal bovine serum (ThermoFischer Scientific) and 1 mM L-glutamine (Silantes). To ensure complete incorporation of labelled amino acids, cells were cultured for at least 8 doublings prior to use. Cells were maintained at 37 °C in a humidified incubator with 5% CO_2_ and were reseeded into fresh culture flasks once at 80% confluency following dissociation with 0.05% (w/v) trypsin-EDTA in PBS. For plating, cell count and viability were automatically determined using a Countess trypan blue assay (ThermoFischer Scientific).

### Stress treatment and TPE-MI labelling

Cells were seeded at 40% confluency into either 25 cm^2^ culture flasks or 6-well plates and cultured overnight. In the case of SILAC-labelled cells, compounds were prepared in fresh heavy-labelled media, and the appropriate vehicle control in an equivalent volume of unlabeled media. Culture media was removed and replaced with treatment media, after which cells were incubated for the duration of the treatment period at 37 °C in a humidified incubator with 5% atmospheric CO_2_. Details of the concentration and duration for each stress treatment are presented in Table 1.

Following treatment, media was removed and replaced with a half-volume of fresh serum free media (either unlabeled or labelled as appropriate) containing TPE-MI to a final concentration of 100 µM. Cells were incubated for 30 min, then immediately washed with 3× excess of PBS containing 10 mM Glutathione to react any remaining TPE-MI. Cells were then washed with PBS, mechanically detached using a cell scraper and centrifuged at 300 *g* for 5 min. For flow cytometry, cells were resuspended in PBS and analyzed using an LSRFortessa flow cytometer (BD Biosciences) as described previously (Chen *et al*, 2017). Cell pellets for proteomics were lysed by resuspension in lysis buffer (150 mM NaCl, 50 mM Tris, pH 8.0, 1% (v/v) IGEPAL CA-630, 0.5 % (w/v) sodium deoxycholate, 0.1% (w/v) sodium dodecyl sulfate) containing cOmplete Mini, EDTA-free Protease Inhibitor Cocktail and 250 U benzonase and incubated on ice for 30 min. Lysate was spun at 20 000 *g* for 30 min to pellet cellular debris and the supernatant collected to a fresh tube. Protein concentration was determined via bicinchoninic acid protein assay (BCA; Thermo Fischer Scientific) using bovine serum albumin as the mass standard. In the case of isotopically labelled cultures, protein from each control and treated sample was combined 1:1 (w/w). Prepared lysates were then precipitated via dilution into a 5-fold excess of ice cold 100% acetone and incubated at –20 °C overnight.

### Sample preparation for mass spectrometry

Samples were centrifuged at 20 000 *g* for 30 min at 4 °C, then the supernatant discarded. Protein pellets were solubilized in 100 µl of 8 M urea in 50 mM triethylammonium bicarbonate (TEAB), and incubated with shaking at 37 °C for 45 min. Proteins were reduced using 10 mM tris(2-carboxyethyl)phosphine, pH 8.0, and alkylated with 10 mM iodoacetamide for 45 min, before being digested with 2 µg trypsin (ThermoFischer Scientific) overnight with shaking at 37 °C. Peptides were then desalted via solid-phase extraction using an Oasis HLB 1 cc Vac Cartridge (catalogue number 186000383, Waters Corp., USA) washed with 1 ml of 80% (v/v) acetonitrile (ACN) containing 0.1% v/v trifluoroacetic acid (TFA), then pre-equilibrated with 2.4 ml of 0.1% (v/v) TFA. Peptides were acidified with formic acid to a final concentration of 1% (v/v), then loaded onto the cartridge and washed with 1.5 ml of 0.1% (v/v) TFA before being eluted in 800 µl of 80% (v/v) ACN containing 0.1% (v/v) TFA. Samples were collected in fresh tubes and lyophilized (VirTis Freeze Dryer, SP Scientific). Peptides were resuspended in 80 µl distilled water and quantified using a BCA assay as above. Peptide aliquots were combined with 5× loading buffer to yield 20 µl containing 2 µg peptides in 2% (v/v) ACN containing 0.05% (v/v) TFA for analysis.

### NanoESI-LC-MS/MS

Samples were analyzed by nanoESI-LC-MS/MS using a Orbitrap Fusion Lumos mass spectrometer (Thermo Scientific) fitted with a nanoflow reversed-phase-HPLC (Ultimate 3000 RSLC, Dionex). The nano-LC system was equipped with an Acclaim Pepmap nano-trap column (Dionex—C18, 100 Å, 75 µm × 2 cm) and an Acclaim Pepmap RSLC analytical column (Dionex—C18, 100 Å, 75 µm × 50 cm). For each LC-MS/MS experiment, 0.6 µg of the peptide mix was loaded onto the enrichment (trap) column at an isocratic flow of 5 µl min^−1^ of 3% CH3CN containing 0.1% (v/v) formic acid for 5 min before the enrichment column was switched in-line with the analytical column. The eluents used for the LC were 0.1% (v/v) formic acid (solvent A) and 100% ACN/0.1% formic acid (v/v) (solvent B). The gradient used (300 nl min^−1^) was from 3–22% B in 90 min, 22–40% B in 10 min and 40–80% B in 5 min then maintained for 5 min before re-equilibration for 8 min at 3% B prior to the next analysis. All spectra were acquired in positive ionization mode with full scan MS from m/z 400–1500 in the FT mode at 120,000 mass resolving power (at m/z 200) after accumulating to a target value 5.00e^5^ with maximum accumulation time of 50 ms. Lockmass of 445.12002 was used. Data-dependent HCD MS/MS of charge states > 1 was performed using a 3 s scan method, at a target value of 1.00e^4^, a maximum accumulation time of 60 ms, a normalized collision energy of 35%, an activation Q of 0.25, and at 15,000 mass resolving power. Dynamic exclusion was used for 45 s.

### Peptide identification and quantitation

Initial identification was carried out using MaxQuant (version 1.6.2.10) against the Swissprot Mus Musculus database (Version: 2016_06; 16794 entries). The search was conducted with 20 ppm MS tolerance, 0.6 Da MS/MS tolerance, with one missed cleavage allowed and match between runs enabled. Variable modifications included methionine oxidation, N-terminal protein acetylation, and N-terminal methionine cleavage while carbamidomethylcysteine was set as a fixed modification. The false discovery rate maximum was set to 0.005% at the peptide identification level (actual was 0.005 for each replicate) and 1% at the protein identification level. All other parameters were left as default.

The change in cysteine peptide abundance following TPE-MI treatment was then determined via custom python scripts (available from DOI: 10.5281/zenodo.6548917 [https://doi.org/10.5281/zenodo.6548917]). The logic was as follows; first, the common contaminant protein keratin was removed. Then, quantified proteins were filtered to those identified by at least two unique peptides, at least one of which contained a cysteine residue. The average peptide abundance for the non-cysteine-containing peptides was then calculated for each protein. These values were used to normalize the cysteine-containing peptides, yielding a corrected cys ratio which accounts for any changes in overall protein abundance due to treatment.

The resultant corrected cysteine and non-cysteine ratios were then scaled using a p-value weighted correction, as described previously (Cox *et al*, 2022). In essence, rather than using the p-value as an arbitrary cut-off, this method scales the mean of biological replicates (*n*=3) according to the relative confidence with which it deviates from the expected value (in this case 0). We then applied a set of thresholds for the cysteine and non-cysteine peptides derived from a control experiment in which both the light-and heavy-labelled were treated with the vehicle control DMSO (Fig. S6). The thresholds were calculated to contain 95% of the control dataset (corresponding to a z-score of 1.96), and datapoints outside these thresholds are considered relevant to the treatment condition. To compare between different treatments, all proteins quantified according to the above criteria (pre thresholding) in all treatments were considered. From the resultant list of proteins, the comparison set contained those for which at least one cysteine-containing peptide exceeded the control threshold in at least one treatment condition. Finally, a summary measure was calculated as the maximum corrected cys ratio per protein in each treatment which was then used for subsequent protein-based comparisons.

### Functional characterization

Physicochemical properties for individual cysteine peptides and proteins of interest were compiled from various databases, including the Protein Data Bank (https://www.ebi.ac.uk/pdbe/), and STRINGdb (v11.0, medium confidence score > 0.4; (Szklarczyk *et al*, 2019)) via Cytoscape v3.9.0 (Shannon *et al*, 2003). Gene ontology annotations for individual proteins were collected from UniProt (https://www.uniprot.org/). Gene ontology enrichment analyses were completed using PantherGOSlim (http://pantherdb.org; (Mi *et al*, 2021)) against the background of all proteins identified in the raw dataset. Significantly enriched terms were filtered according to p < 0.05, and the most specific terms from each hierarchically redundant family are presented. Connected clusters were detected in the protein-protein interaction map using the Girvan-Newman fast greedy algorithm (Newman & Girvan, 2004) for community detection, as implemented by the cytoscape Glay plugin (Su *et al*, 2010). To compare potential sources of correlation among individual proteins, a series of feature bins were considered; namely, community cluster, KEGG pathways, protein-protein interactions, and complex memberships. The correlation strength between individual proteins was determined as the absolute Spearman’s correlation coefficient (R_s_) for individual protein pairs, and each pair was then binned according to whether one (outside) or both (inside) proteins identified with the feature of interest. Pairs for which neither protein identified with the feature of interest were discarded. For each protein, the mean correlation inside and outside the feature bin was then calculated. Finally, for features associated with at least 3 proteins, the mean correlation across all feature proteins was compared.

### Comparison to Sui *et al* dataset

Summary data from Sui et al. (Sui *et al*, 2020) was downloaded from the supplementary information available online 10.1073/pnas.1912897117 [https://doi.org/10.1073/pnas.1912897117]. Datasets for treatment conditions common to both studies were collected (MG132, VER155008 and novobiocin), and the pellet-based solubility ratio (Sui dataset) was compared with the maximum corrected cysteine ratio (TPE-MI dataset) for individual proteins. Proteins altered by treatment in both datasets were collected, and in cases where more than three proteins passed this filter their correlation was assessed via linear regression.

### Data availability and statistical analysis

The mass spectrometry proteomics data have been deposited to the ProteomeXchange Consortium via the PRIDE63 partner repository with the data set identifier PXD033152. Preprocessed datasets for the proteomics and flow cytometry are also available from DOI 10.5281/zenodo.6439170 [https://doi.org/10.5281/zenodo.6439170].

Statistical analyses were performed using either the scipy module in python (Virtanen *et al*, 2020) or GraphPad Prism (v 8.4.3). The exact *p*-values and statistical details are provided in Supplementary Dataset 1. All analysis scripts are available from DOI: 10.5281/zenodo.6548917 [https://doi.org/10.5281/zenodo.6548917].

## Acknowledgements

We thank Dr. Yuning Hong (La Trobe University) for providing the TPE-MI reagent, and the Bio21 Melbourne Mass Spectrometry and Proteomics facility for sample processing.

## Author contributions

D.C., G.E.R., and D.M.H. designed the research; D.C. and A.O. performed the research and analyzed the data; and D.C. wrote the manuscript, and all authors revised and approved the final version.

## Supplementary Figures

**Figure S1:**
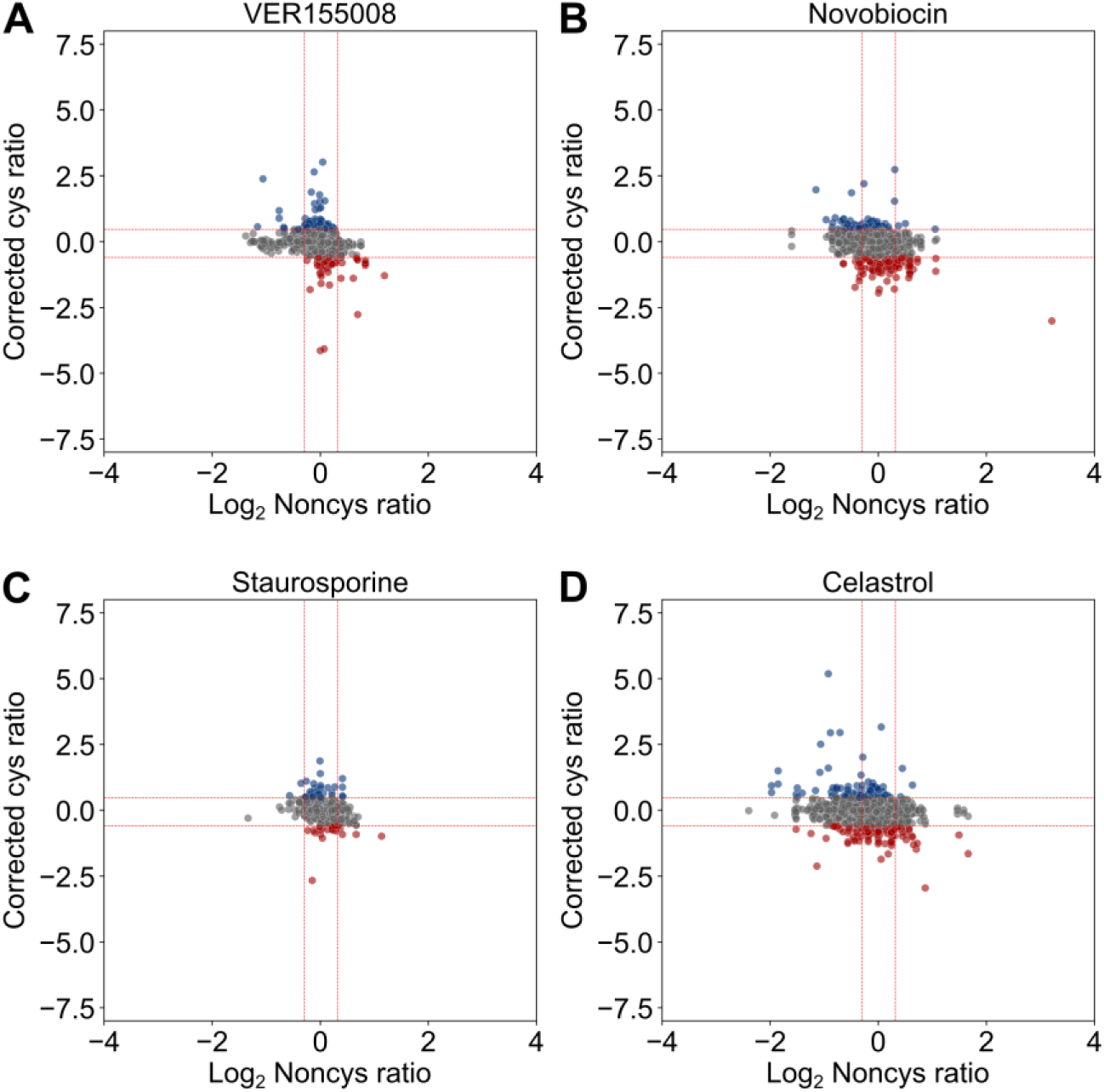
Summary of cysteine thiol reactivity changes associated with individual pharmacological stimuli. Scatterplots for processed noncysteine-and cysteine-peptide ratios in the presence of (A) VER155008, (B) novobiocin, (C) staurosporine, and (D) celastrol. Thresholds (red dotted line) determined based on control dataset, outside which cysteine-containing peptides are considered to be more exposed (red) or more protected (blue) due to treatment.

**Figure S2:**
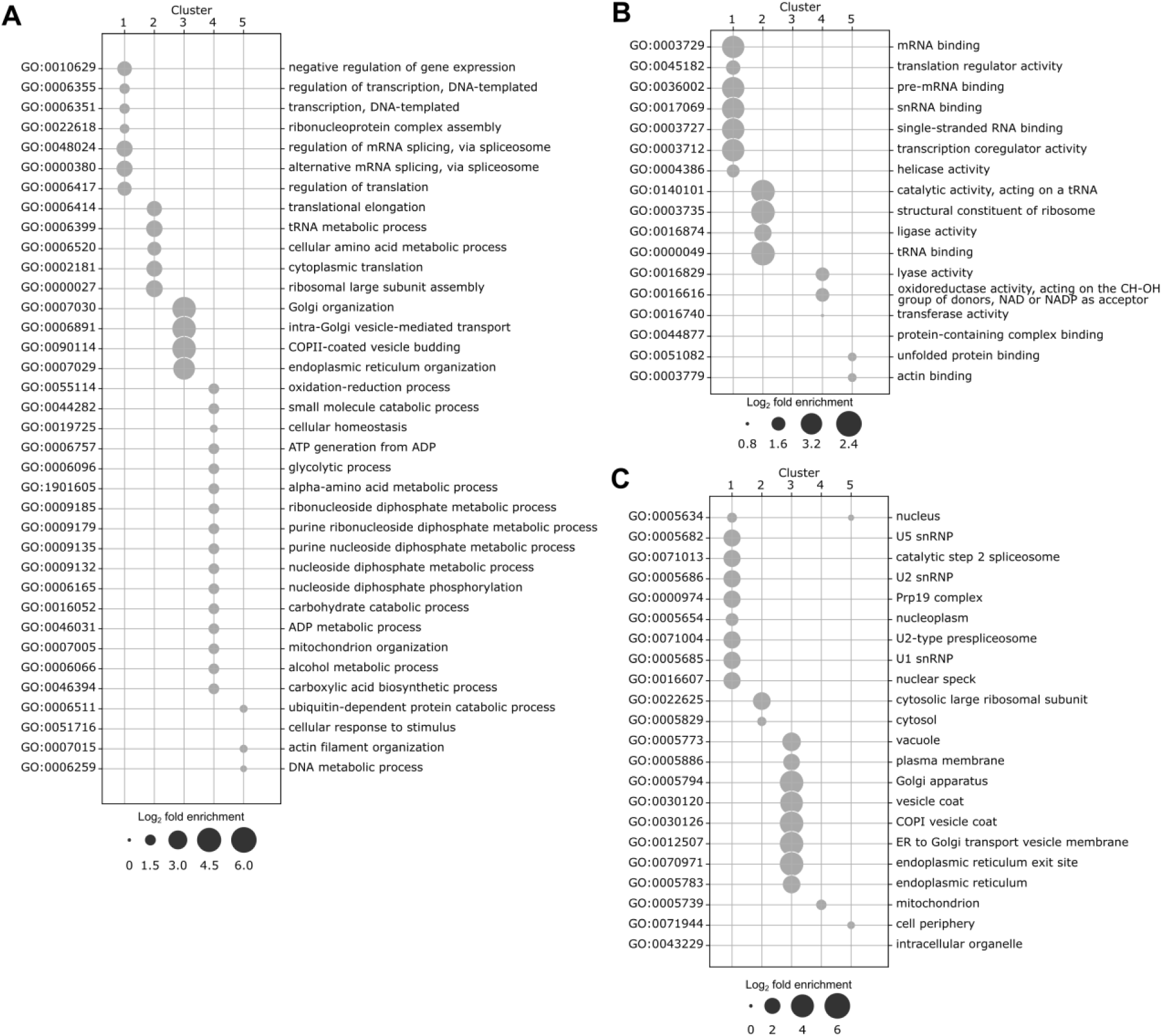
Gene ontology enrichment for each protein-protein interaction cluster. Significantly enriched gene ontology terms (p < 0.05) for proteins found in functional clusters 1 -5. Independent search results are shown for (A) biological process, (B) molecular function or (C) cellular component, and enrichment terms were filtered to minimize hierarchical redundancy among ontology families (PantherGOSlim v 16.0).

**Figure S3:**
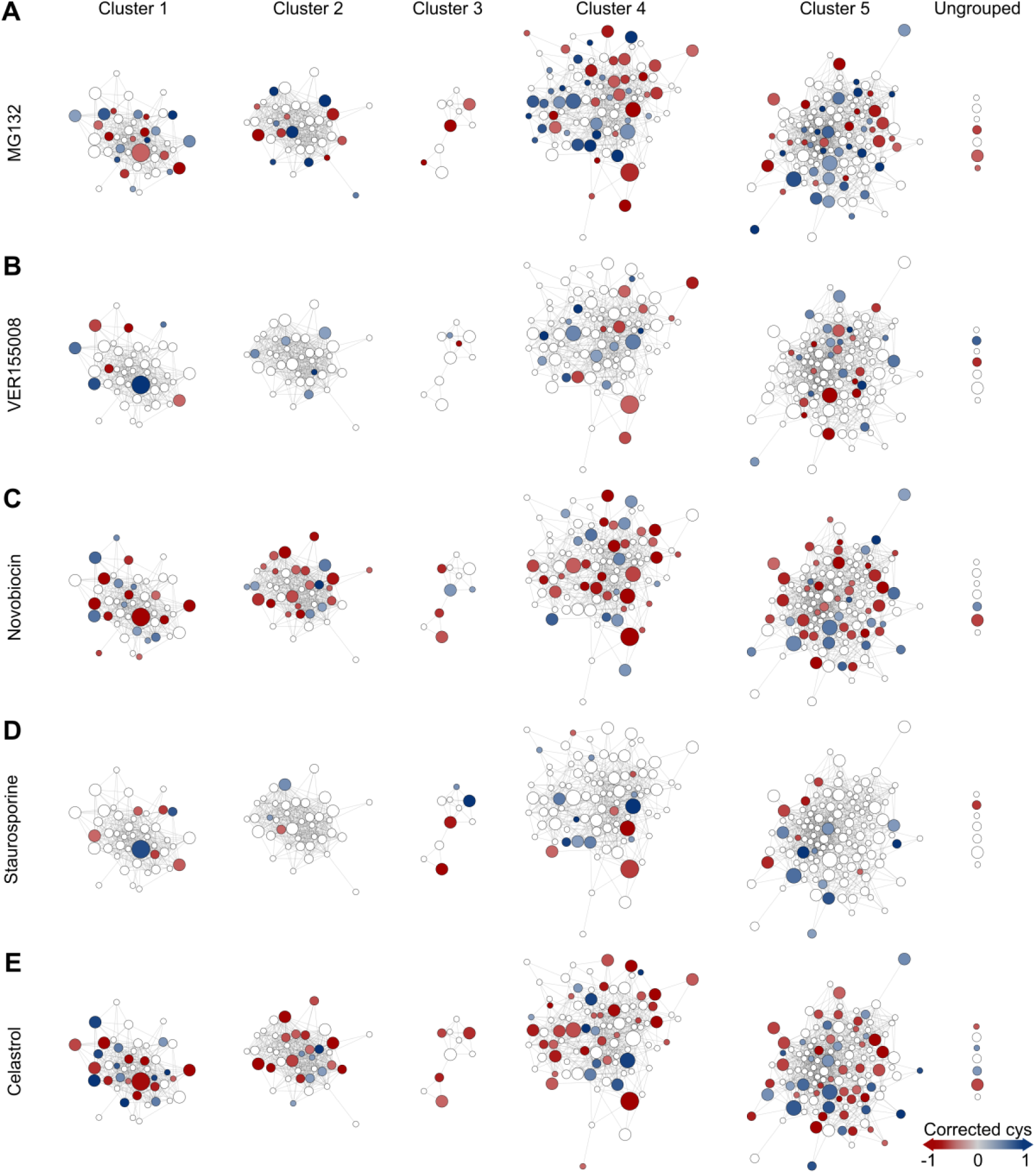
Per-protein changes in cysteine reactivity are heterogeneous. Clustered protein interaction network for comparison proteins. Protein nodes are sized according to degree of commonality across treatments and colored according to maximum corrected cysteine thiol ratio change due to individual treatments. Nodes were arranged organically following clustering with Girvan-Newman community detection algorithm, and edges (lines) connect proteins with known interactions within each cluster (STRINGdb v 11.0, medium confidence score > 0.4).

**Figure S4:**
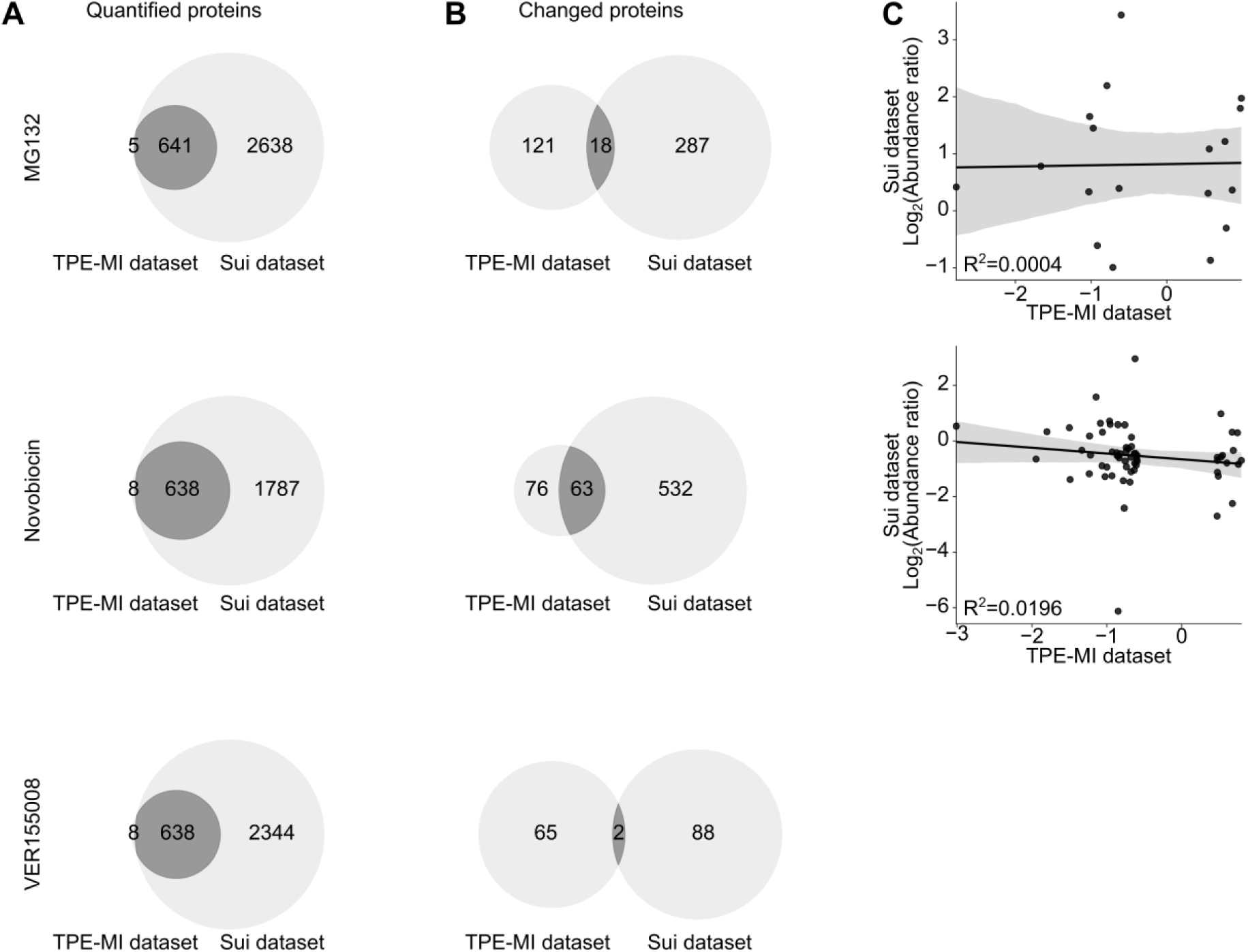
Correlation between cysteine thiol reactivity and solubility changes measured in MG132, novobiocin or VER155008. The overlap between proteins (A) identified or (B) significantly changed according to the Sui *et al*. dataset *(Sui* et al, *2020)* compared with the comparison set of proteins (TPE-MI dataset) quantified in the present study. (C) Correlation between the Sui *et al*. and TPE-MI datasets in which more than three common proteins were identified as having undergone a significant change due to treatment. Confidence interval derived via automatic bootstrap estimation of the linear regression.

**Figure S5:**
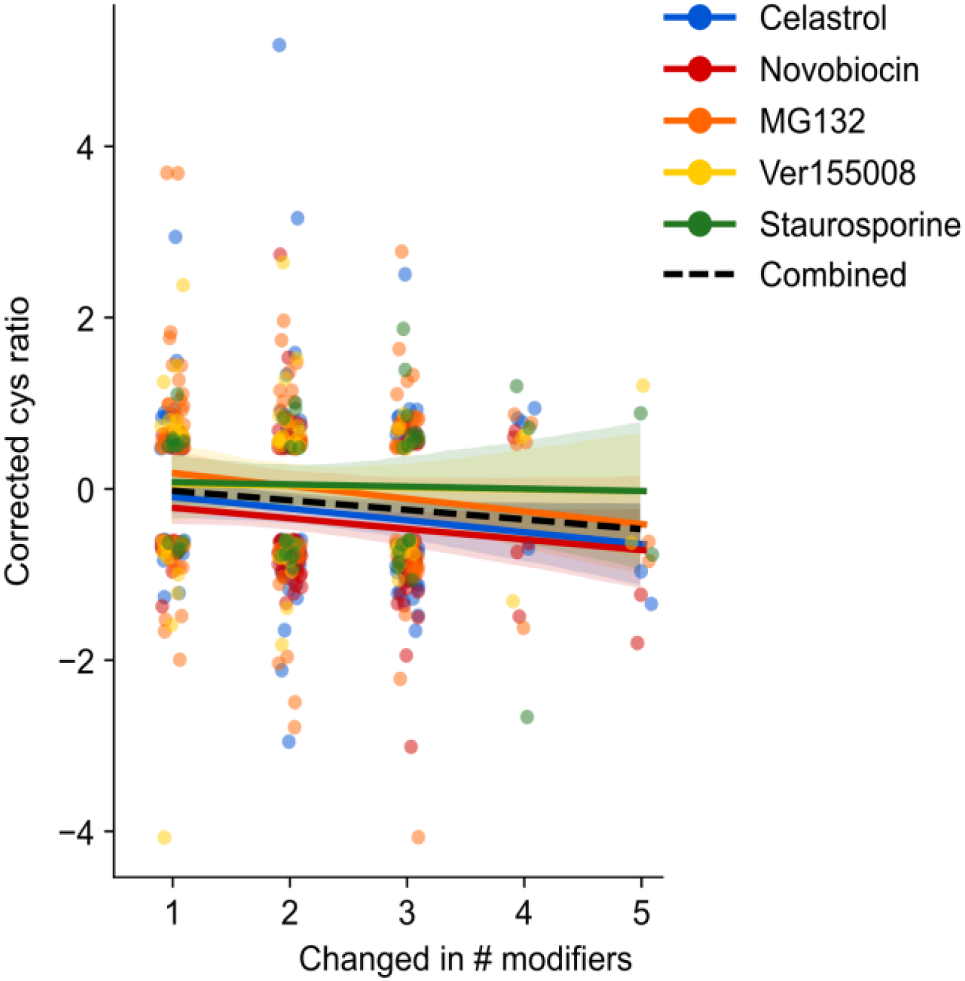
Correlation between conservation degree and maximum cysteine thiol reactivity change per protein. Comparison proteins are binned according to their degree of conservation across treatments. Correlation and corresponding confidence intervals were derived via automatic bootstrap estimation of the linear regression for either individual treatments (colored samples) or the comparison dataset as a whole (black dashed line).

**Figure S6:**
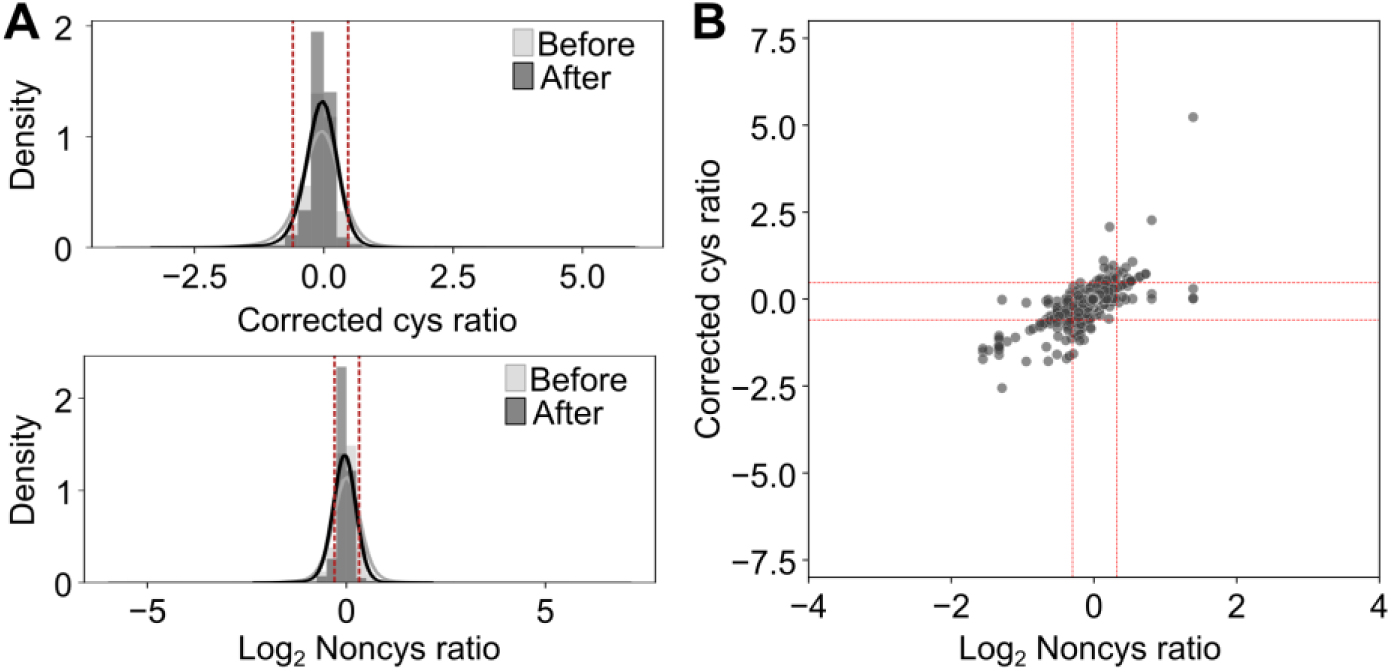
Threshold derivation from control dataset. Histograms for the control vs control dataset of (A) cysteine and (B) noncysteine-containing peptide ratios before and after p-value scaling. The z-score was calculated for each peptide, and thresholds set according to values at which the z-score surpassed 1.96, such that 95% of the control data is contained within the thresholds. (C) The resultant thresholded scatterplot for the control dataset.

